# Ancient DNA from Protohistoric Period Cambodia indicates that South Asians admixed with local populations as early as 1^st^-3^rd^ centuries CE

**DOI:** 10.1101/2022.06.30.498315

**Authors:** Piya Changmai, Ron Pinhasi, Michael Pietrusewsky, Miriam T. Stark, Rona Michi Ikehara-Quebral, David Reich, Pavel Flegontov

## Abstract

Indian cultural influence is remarkable in present-day Mainland Southeast Asia (MSEA), and it may have stimulated early state formation in the region. Various present-day populations in MSEA harbor a low level of South Asian ancestry, but previous studies failed to detect such ancestry in any ancient individual from MSEA. In this study, we discovered a substantial level of South Asian admixture (ca. 40% – 50%) in a Protohistoric individual from the Vat Komnou cemetery at the Angkor Borei site in Cambodia. The location and direct radiocarbon dating result on the human bone (95% confidence interval is 78 – 234 calCE) indicate that this individual lived during the early period of Funan, one of the earliest states in MSEA, which shows that the South Asian gene flow to Cambodia started about a millennium earlier than indicated by previous published results of genetic dating relying on present-day populations. Plausible proxies for the South Asian ancestry source in this individual are present-day populations in Southern India, and the individual shares more genetic drift with present-day Cambodians than with most present-day East and Southeast Asian populations.

## Introduction

The high ethnolinguistic diversity of Mainland Southeast Asia (MSEA) reflects the complex population history of this region^1^. Anatomically modern humans arrived in MSEA approximately 50,000 years ago^2^. The early to mid-Holocene hunter-gatherers associated with the Hoabinhian archaeological tradition (with genomic data available for individuals from Laos and Malaysia dated to ca. 8,000 and 4,500 years before present, respectively) were modelled as a deeply diverged East Eurasian lineage, related to present-day Andamanese and to MSEA foragers such as Jehai^3^. During the Neolithic period (starting at ∼4,500 years before present, BP), ancient MSEA populations exhibited an admixed genetic profile between the deeply diverged East Eurasian lineage and an East Asian^3,4^. The same genetic mixture is typical for some present-day Austroasiatic-language speaking groups, such as the Mlabri^4^. The genetic structure of Bronze Age individuals from Northern Vietnam (dated to ∼2,000 BP) suggests an additional wave of migration from Southern China^3,4^.

Key debates over the origins of early states in MSEA have hinged on the nature of contact with South Asia for more than 50 years (e.g., Cœdès 1968^5^). Archaeological evidence revealed that the cultural interaction between Indian and local MSEA people began by the 4^th^ century BCE, creating synergies that influenced the formation of early states in MSEA^6,7^. Various present-day MSEA populations, especially those highly influenced by Indian culture, harbor a low level of South Asian admixture: for instance, Bamar, Cham, Khmer, Malay, Mon, and Thai^8-10^. Using methods based on autosomal haplotypes, this genetic admixture was dated to around the 14^th^ c. CE (1194 CE – 1502 CE) in a Khmer group from Cambodia^11^). In a later study, the admixture date for the Khmer group from Cambodia was confirmed (12^th^-13^th^ cc. CE), and admixture date estimates for a wider range of present-day SEA groups were obtained: they lie between the 16^th^ c. CE for Bamar and the 4^th^ cc. CE for Giarai (Jarai) from Vietnam^9,10^. Previous ancient DNA studies, although of very limited extent due to poor ancient DNA preservation in tropical climate, did not detect South Asian ancestry in any ancient MSEA individuals^3,4^.

Here we report a finding of substantial South Asian ancestry in a Protohistoric Period male child from the Vat Komnou cemetery at the Angkor Borei site on the western edge of Cambodia’s Mekong Delta. The walled and moated Angkor Borei site was first occupied in the middle of the first millennium BCE and is one of the earliest dated urban sites in the Mekong Delta in either Cambodia or Vietnam^12^. In addition to brick architectural monuments, associated moats, and ponds, the Vat Komnou mortuary assemblage includes human burials, beads, ceramics, multiple pig skulls, and other faunal remains^13,14^. The Vat Komnou cemetery is one of the largest archaeological skeletal samples analyzed to date from Cambodia.

Genetic data for the male child from the Vat Komnou cemetery (skeletal code AB M-40) were first reported by Lipson et al. (2018)^4^: 0.047x coverage on the 1240K SNP panel used for enrichment of ancient DNA^15,16^, and we increased the coverage to 0.061x and generated variant calls at 64,103 1240K sites (as compared to 54,221 sites in Lipson *et al*.^4^). A direct radiocarbon date on the human bone (95% confidence interval is 78 – 234 calCE) was also obtained for the individual AB M-40 by Lipson *et al*. (2018)^4^.

## Results

### Genetic overview of ancient Southeast Asians

The location of the ancient individual from Cambodia (AB M-40 or I1680) is illustrated in Fig. 1. We explored genetic ancestry of this individual and other ancient individuals from Southeast Asia for whom published genetic data of acceptable quality were available (i.e., pseudo-haploid genotypes are available for more than 50,000 1240K sites). We computed principal components (PCs) using present-day populations from Europe, from East, Southeast, South, and Central Asia, the Andaman Islands, and Siberia, and projected 28 ancient individuals (from McColl et al., 2018^3^, Lipson et al., 2018^4^, and this study) onto these PCs. The PC1 vs. PC2 plot (Fig. 2, Suppl. Fig. 1) reveals three clusters of individuals: East and Southeast Asian (ESEA), European (EUR), and Andamanese Negrito. South Asian (SAS) populations form a cline between the European and Andamanese Negrito clusters, with populations from Pakistan and Northern India lying closer to the European cluster, in line with previous studies on the population genetics of South Asia^17-19^. The positions of populations speaking languages of the Munda branch of the Austroasiatic language family, such as Birhor and Kharia deviate from the South Asian cline, and this is not unexpected since they were shown to harbor Southeast Asian admixture^18,20^. Kusunda, a group from Nepal, and Riang, another Austroasiatic-speaking group from India, fall close to the East and Southeast Asian cluster. These populations harbor a relatively high proportion of an ancestry component maximized in ESEA populations according to our *ADMIXTURE* analysis (Fig. 3). Central Asian and Siberian populations are distributed along PC1 between the SAS – EUR cline and the ESEA cluster, in line with many previous studies, such as Jeong et al., (2019)^21^. Most ancient individuals dated to 2500 BCE – 1950 CE from Southeast Asia fall within the ESEA cluster, except for the individual AB M-40/I1680 from the Vat Komnou site who deviates from the ESEA cluster towards the South Asian cline. The *ADMIXTURE*^22^ analysis also indicates that this individual stands out among Southeast Asians since his genome harbors a high proportion of both “blue” and “grey” ancestry components, maximized in European and Andamanese populations, respectively. Both components are also prominent in South Asian populations (Fig. 3). Present-day Cambodians share more drift with the Protohistoric individual AB M-40/I1680 from Cambodia than most present-day ESEA populations as inferred by “outgroup” *f*_*3*_-statistics *f*_*3*_(Mbuti; AB M-40/I1680, X) (Fig. 4). To sum up, these PCA and *ADMIXTURE* results suggest that substantial South Asian admixture is present in the individual AB M-40/I1680, which sets him apart from 27 other Southeast Asians for whom genetic data of acceptable quality was reported in the literature^3,4^.

**Fig. 1.**
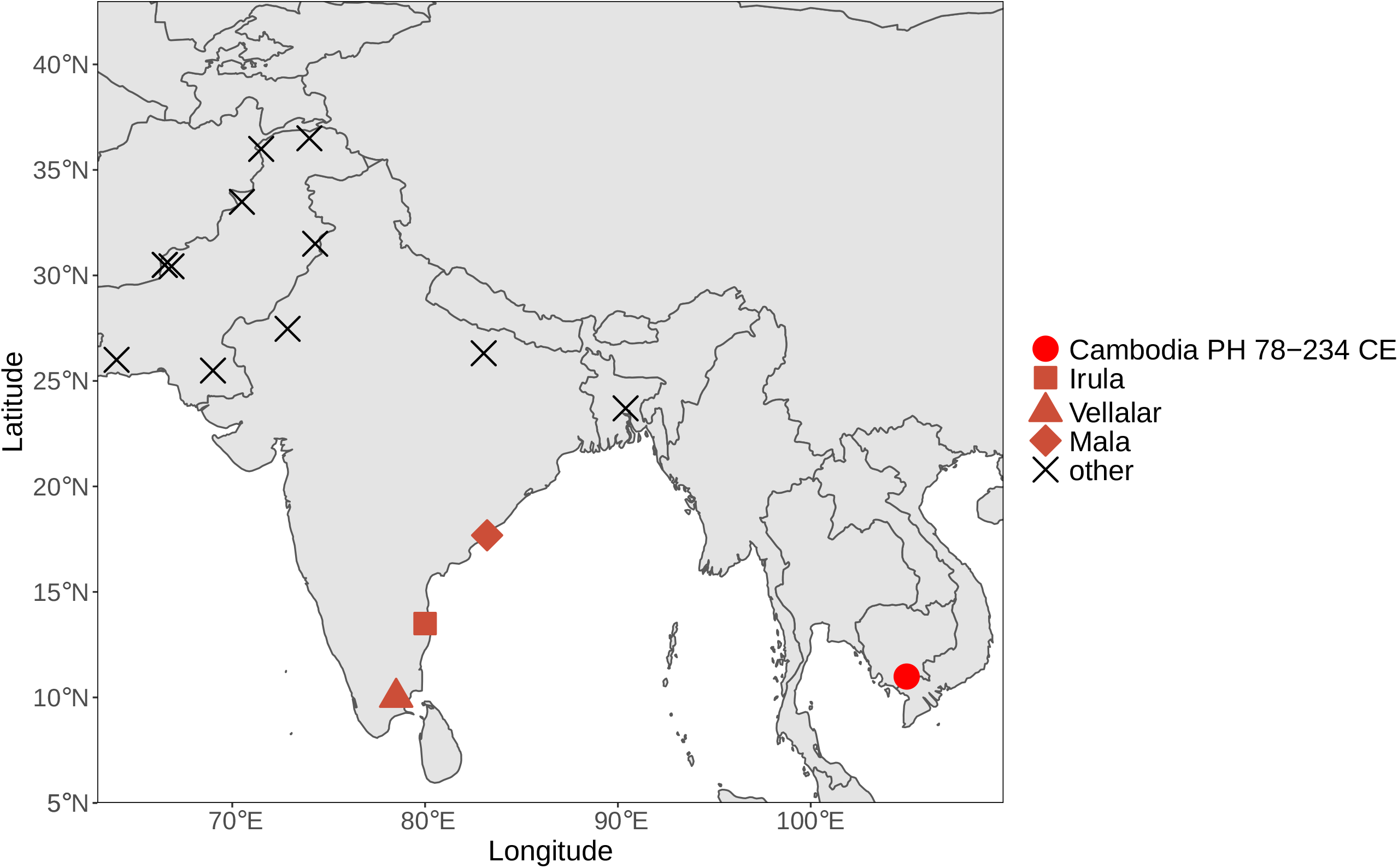
Geographic locations of the ancient individual AB M-40/I1680 sequenced in this study (the circle) and of the present-day South Asian surrogates used in *qpAdm* analysis (other marker shapes). The South Asian surrogates which are not included in any plausible *qpAdm* model are labeled with black cross markers.

**Fig. 2.**
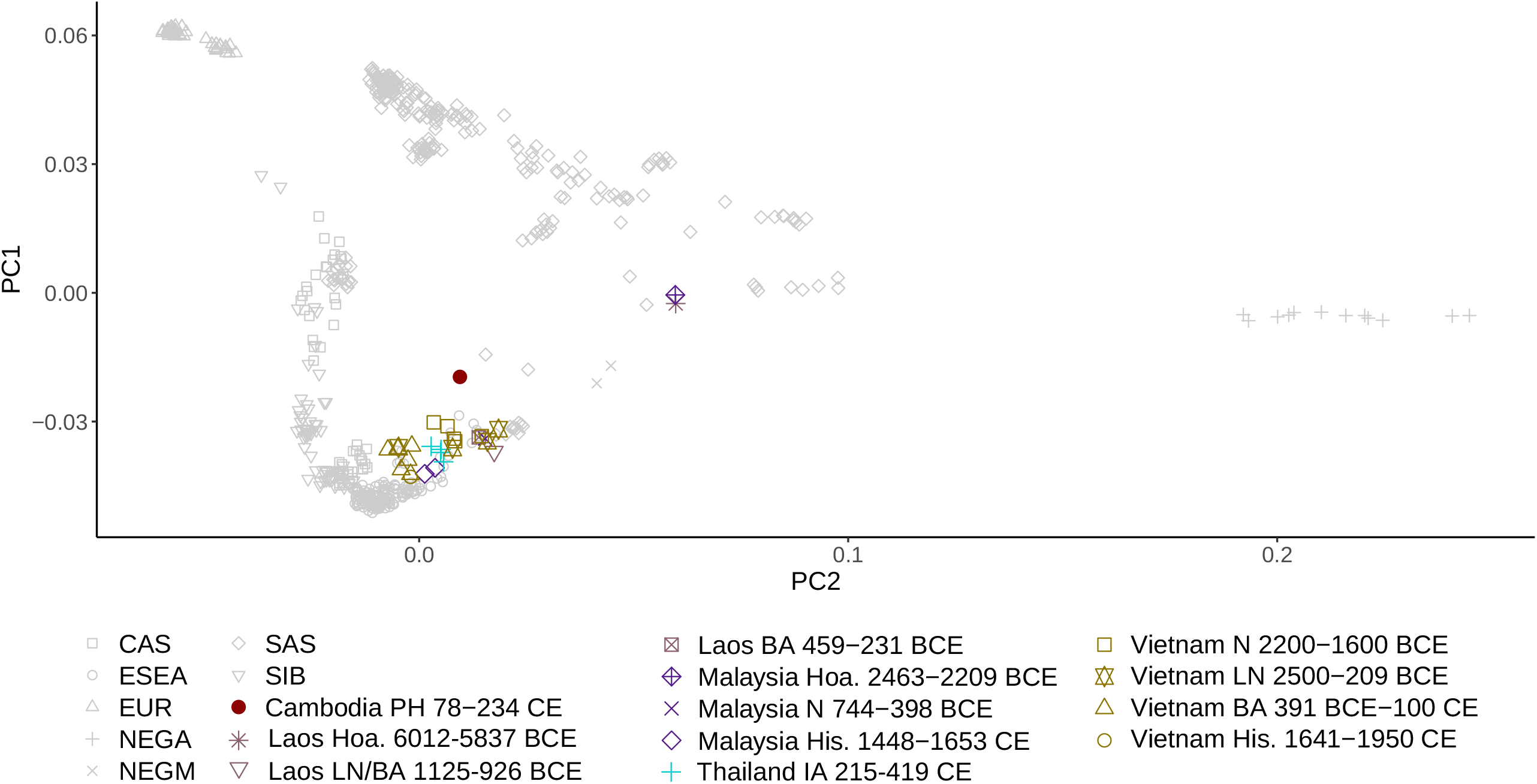
Principal component analysis (PCA). Ancient Southeast Asian individuals were projected on the principal components calculated using present-day Eurasian populations, and the first two components (PC1 and PC2) are shown. Abbreviations for meta-populations are as follows: CAS, present-day Central Asians; ESEA, present-day East and Southeast Asians; EUR, present-day Europeans; NEGA, present-day Andamanese Negritos; NEGM; present-day Mainland Southeast Asian Negritos; SAS, present-day South Asians; SIB, present-day Siberians; Hoa., Hoabinhian culture; N, Neolithic; LN, Late Neolithic; PH, Protohistoric period; BA, Bronze Age; His., Historical period.

**Fig. 3.**
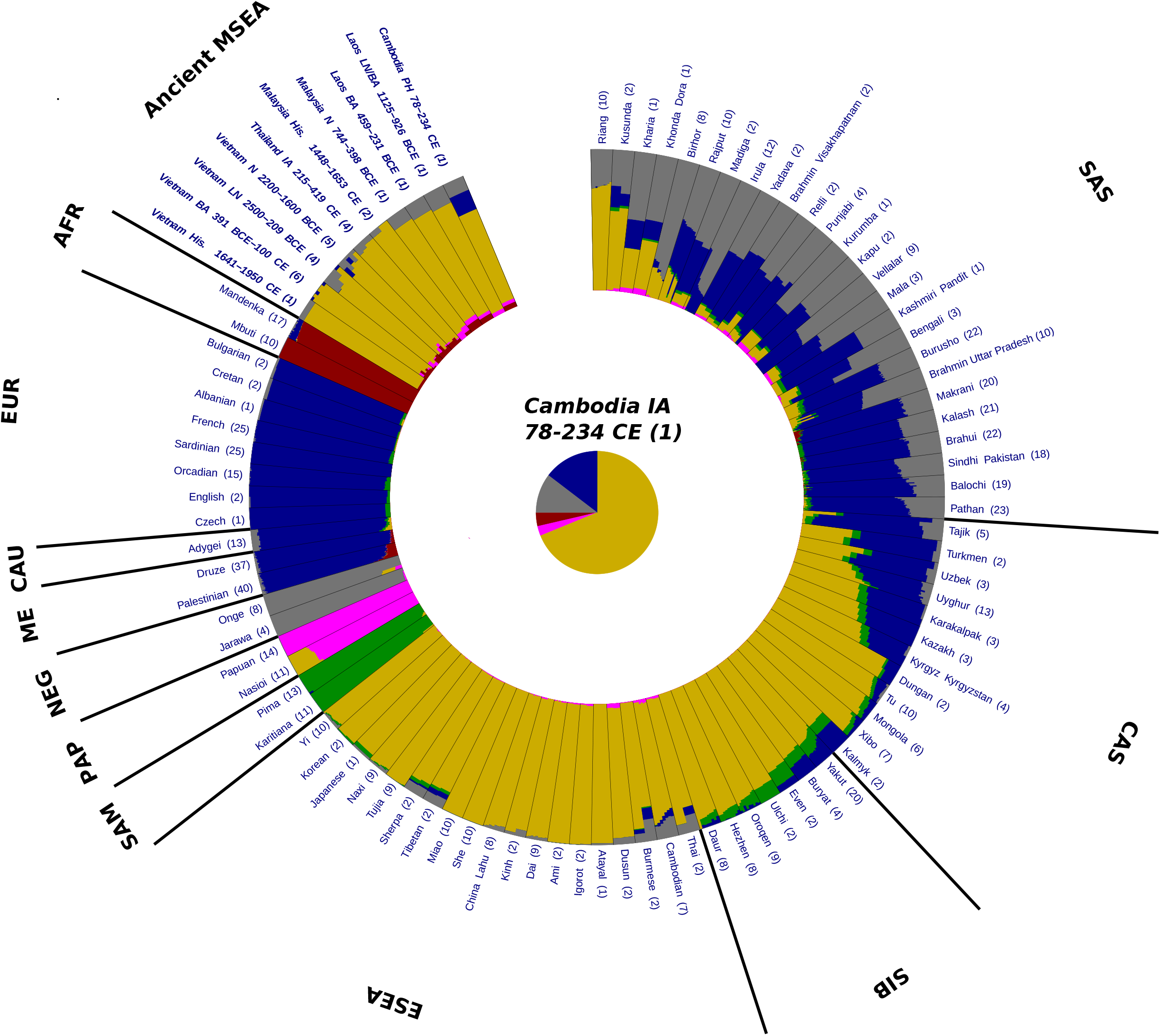
An *ADMIXTURE* analysis plot showing results for 6 hypothetical ancestral groups (*K*=6). Abbreviations for meta-populations are as follows: Ancient MSEA, Ancient Mainland Southeast Asians; AFR, present-day Africans; EUR, present-day Europeans; CAU, present-day Caucasians; ME, present-day Middle Easterners; NEG, present-day Andamanese Negritos; PAP, present-day Papuans; SAM, present-day Native Meso- and South Americans; ESEA, present-day East and Southeast Asians; SIB, present-day Siberians; CAS, present-day Central Asians; SAS, present-day South Asians; N, Neolithic; LN, Late Neolithic; PH, Protohistoric period; BA, Bronze Age; His., Historical period. The number of individuals for each population is indicated in brackets after the population name.

**Fig. 3.**
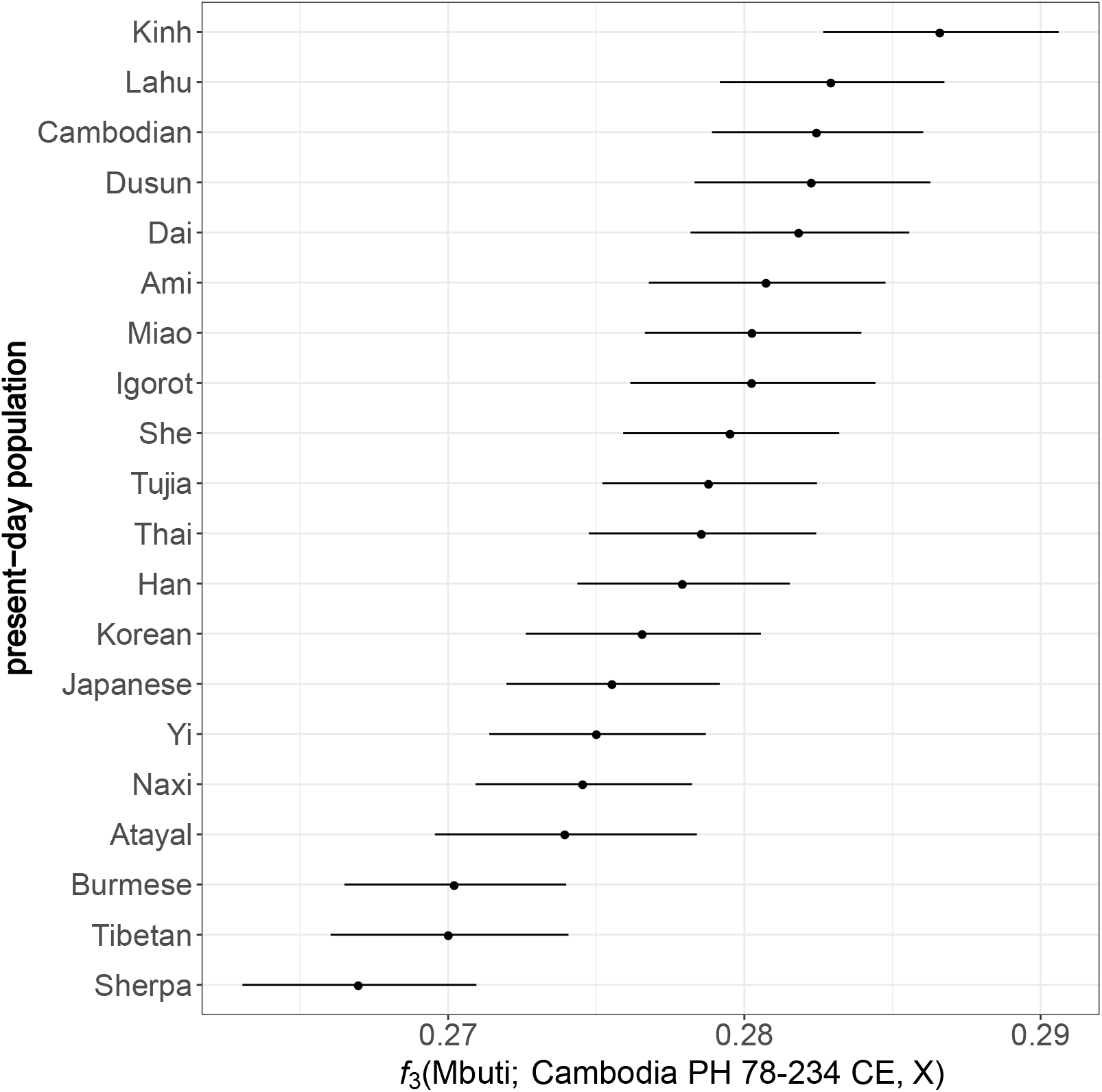
Genetic affinity between the ancient individual AB M-40/I1680 and present-day East and Southeast Asian groups. Genetic drift sharing was estimated using outgroup *f*_*3*_-statistics in the form of *f*_*3*_(Mbuti; AB M-40/I1680, X), where X is a present-day East and Southeast Asian group.

### South Asian ancestry in ancient Southeast Asians

We explored further the signal of South Asian admixture in ancient Southeast Asians by inspecting a scatterplot of outgroup *f*_*3*_-statistics, *f*_*3*_(Mbuti; South Asian, X) and *f*_*3*_(Mbuti; East Asian, X), where X are ancient individuals from Southeast Asia, present-day ESEA populations, Tibeto-Burman-speaking groups and Kusunda from Nepal, and Austroasiatic-speaking populations from India (Fig. 5). We used the Brahmin from Uttar Pradesh^23^ and Dai^24^ populations as South Asian and East Asian surrogates, respectively. Most present-day and ancient groups from ESEA demonstrate a linear relationship of genetic drift sharing with the South Asian and East Asian surrogates. *f*_*3*_-statistics for various groups, namely the ancient individual AB M-40/I1680 from Cambodia, Tibeto-Burman-speaking populations (Tibetan and Sherpa) and Kusunda from Nepal, and Austroasiatic-speaking groups from India (Birhor, Kharia, and Riang), do not conform to this trend line (Fig. 5). The shift indicates excessive shared genetic drift with the South Asian surrogate in these populations compared to most ESEA groups.

**Fig. 5.**
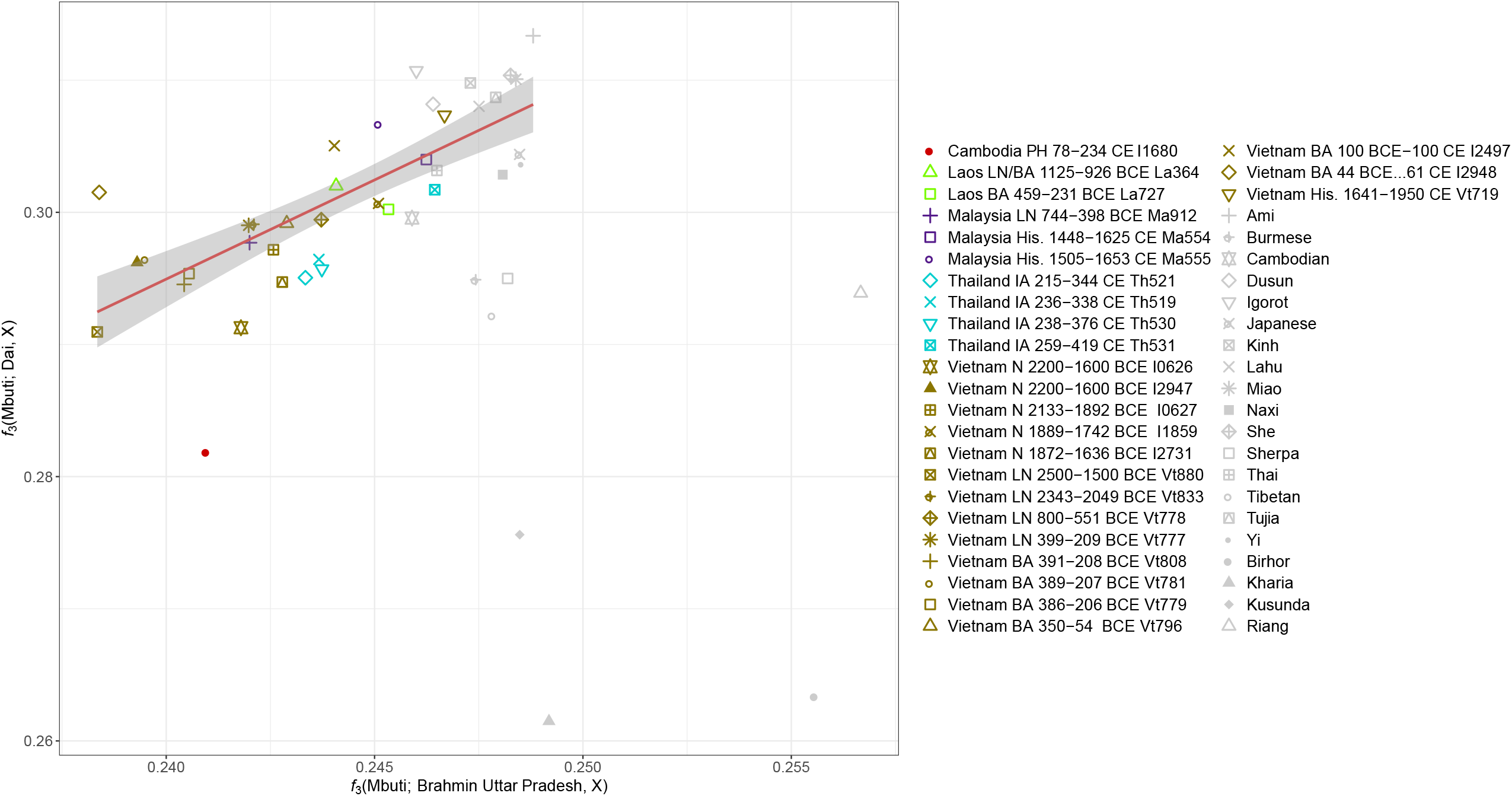
A biplot of outgroup *f*_3_-statistics. The plot of *f*_3_(Mbuti; Brahmin Uttar Pradesh, X) vs. *f*_3_(Mbuti, Dai, X), where X is an ancient individual from Southeast Asia or present-day populations from East, Southeast, or South Asia. The plot demonstrates shared genetic drift between population X and Brahmin from Uttar Pradesh (a SAS population) or Dai (an ESEA population). The trend line illustrates the ratio of genetic drift sharing between most ESEA populations and the SAS and ESEA surrogate. Abbreviations: N, Neolithic; LN, Late Neolithic; PH, Protohistoric period; BA, Bronze Age; His., Historical period.

Notably, South Asian admixture in Bamar, Tibetans, Sherpa, and Austroasiatic-speaking populations from India has been reported in previous studies^8,18,20,25,26^.

We formally tested for South Asian admixture in the ancient individual AB M-40/I1680 from Cambodia using a *qpAdm*^27^ protocol with a “rotating” set of reference or “right” populations^28^. We first performed pairwise *qpWave* tests to check if the target ancient individual is cladal (genetically continuous) with a single reference population (see Materials and Methods for details). None of these pairwise *qpWave* models were plausible (Suppl. Table 1). Next, we tested all possible pairs of ancestry sources (78 pairs) from the set of reference populations (see Materials and Methods), and the genome the individual of interest can be modelled as a two-way mixture of East Asian and South Asian populations (Fig. 6, Suppl. Table 1). Among plausible models (those with *p*-value is higher than 0.05 and with inferred admixture proportions ± 2 standard errors between 0 and 1 for all ancestry components) for the Protohistoric Cambodian individual AB M-40/I1680, Ami is the only East Asian surrogate, and Irula, Mala, and Vellalar are the only non-East Asian surrogates that result in plausible models (Fig. 6, Suppl. Table 1). Remarkably, all these surrogates are from Southern India (Fig. 1), and the fraction of Southern Indian ancestry was estimated at 42% to 49% across all plausible *qpAdm* models (Fig. 6, Suppl. Table 1). The fact that *qpAdm* models including European, Middle Eastern, Caucasian, or even Northern Indian ancestry sources are rejected suggests that contamination with modern DNA is an unlikely explanation for our results. Moreover, the libraries passed routine ancient DNA authenticity checks such as damage rate at a terminal read nucleotide: a damage rate below 0.03 is considered problematic^4^. See Suppl. Table 2 for detailed statistics for the merged data and for each library.

**Fig. 6.**
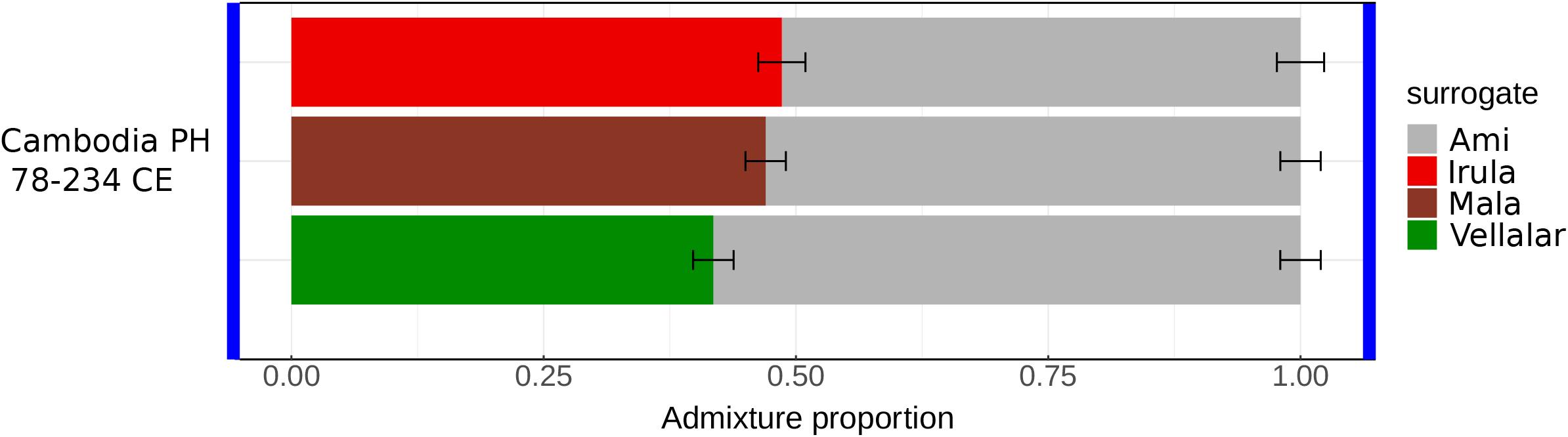
Admixture proportions estimated by *qpAdm*. The plot illustrates all plausible *qpAdm* models inferred using the rotating strategy (see details in Materials and Methods). All the plausible models consist of two surrogates: 1) an East and Southeast Asian surrogate (Ami, on the right-hand side of each bar) and 2) a South Asian surrogate (Irula, Mala, and Vellalar on the left-hand side of each bar). Results for all *qpWave* and *qpAdm* models are available in Suppl. Table 1.

## Discussion

We report new archaeogenomic data from a radiocarbon-dated individual AB M-40/I1680 from Cambodia (95% confidence interval for the date is 78 – 234 calCE) that serves as a key data point for dating the beginning of the South Asian gene flow into Southeast Asia. This individual was previously analyzed by Lipson *et al*. (2018)^4^, but we generated additional data to increase the sensitivity of the analyses in the current study. South Asians were not considered as a potential ancestry source for MSEA groups in the original study, which explains the discrepancy between our results and those by Lipson *et al*. (2018)^4^. For instance, PCA in Fig. 1 of that study lacks any West Eurasian groups, and another PCA in Fig. S1 of that study lacks any South Asian groups except for the Andamanese^4^. Nevertheless, according to the latter PCA^4^ the individual AB M-40/I1680 is shifted towards Europeans and Andamanese as compared to all present-day and most ancient MSEA individuals.

In addition to our genetic results, other anthropological results on the individual AB M-40/I1680 are compatible with his admixed genetic profile. Slight expression of Carabelli’s cusp was observed in the maxillary permanent right and left first molars and left second molar of this individual, and some of the highest frequencies of this dental trait have been reported for the present-day inhabitants of Western Eurasia and India^29^. Strontium isotopic analysis of this individual suggests limited evidence of human mobility^30^.

South Asian admixture is present in various present-day MSEA populations, especially those heavily influenced by Indian culture^8-10^. However, previous aDNA studies did not detect South Asian admixture in ancient individuals across MSEA, although due to the immense technical difficulties genomic data (of any quality) were generated for just 21 individuals from Cambodia, Laos, Malaysia, Thailand, and Vietnam dated to the 5^th^ c. BCE and later^3,4^. As suggested by archaeological evidence^6,7^, we expect to find traces of interactions with South Asia starting in the 4^th^ c. BCE.

An earlier study estimated the proportion of South Asian ancestry in present-day Cambodians at ca. 9%^10^. Notably, the proportion of South Asian ancestry in the ancient individual from Cambodia (point estimates ranging from 42% to 49% across all plausible *qpAdm* models) is considerably higher than in the present-day majority population of the country and in any present-day MSEA population investigated in the literature (see an overview in Changmai et al., 2022^10^). Since we are dealing with a single individual only, it is hard to estimate the intensity of the gene flow and to extrapolate this estimate to the general Protohistoric population in the territory of Cambodia.

The location and date (1^st^-3^rd^ c. CE) of the individual AB M-40/I1680 fit the early period of Funan, an early state in the territories of Cambodia and Vietnam. Chinese written sources documented that the Funan dynasty was established by an Indian Brahmin named Kaundinya and a local princess named Soma^31^. Archaeological evidence from glass and stone beads recovered from the Mekong Delta and peninsular Thailand^32^ and archaeobotanical remains^33^ suggests the possibility of multi-ethnic residence in areas of Protohistoric MSEA whose populations engaged in maritime trade (e.g., Bellina 2014^34^). Collectively, these data suggest some level of Indian cultural influence in the Mekong Delta in the 1^st^-3^rd^ c. CE. The only plausible South Asian genetic sources for the individual from Funan, inferred by the *qpAdm* method, are populations from Southern India. Even though the results suggest that South Indian populations are by far the most plausible surrogates for this individual, we caution that the actual ancient sources possibly had a genetic profile different to the present-day South Asian populations we used for the *qpAdm* analysis.

Consensus now holds that the early first-millennium Mekong Delta kingdoms associated with the Chinese-named Funan were the predecessor states for the Angkorian empire, but what language was spoken before the 7^th^ century CE (when the earliest inscription in the Delta appeared, at Angkor Borei^35^) remains a mystery. The term “Funan” is based on the modern Chinese pronunciation. Whether the local pronunciation was “biunâm”, resembling the old Khmer word “bnam”, meaning “mountain,” remains an unresolved question^5^. Continuities in architectural style, imagery, and settlement forms from the Protohistoric through later Angkorian period suggest that some residents of Funan spoke old Khmer. Outgroup *f*_*3*_-statistics (Fig. 4) support this point as they show that Cambodians and Vietnamese are present-day groups sharing the highest amount of genetic drift with the ancient individual AB M-40/I1680 from Funan.

Changmai et al., (2022)^10^ estimated the date of South Asian admixture events in various present-day MSEA populations between the 4^th^ and 16^th^ cc. CE. The inferred dates of South Asian admixture in present-day Cambodians and in a Khmer group from Thailand are 771 – 808 BP and 1218 – 1291 BP (95% confidence intervals), respectively^10^, but the date of the ancient individual AB M-40/I1680 is much older (1716 – 1872 calBP, 95% confidence interval). Our results suggest that some South Asians migrated to MSEA and intermarried with local people before or at the early stage of state formation. These South Asians may have influenced the expansion of Indian culture and the establishment of Indian-style states (also known as Indianized states^5^). aDNA data from MSEA are scarce, so further data collection is necessary to trace the incorporation of South Asian ancestry into the populations of this region.

## Materials and methods

### Archaeological context, ancient sample preparation, and dataset compilation

Archaeological field investigations at the Vat Komnou cemetery (Angkor Borei, Cambodia) were undertaken by the Lower Mekong Archaeological Project (LOMAP) in 1999 and 2000 as a collaboration between the University of Hawai‘i-Mānoa (USA), the Ministry of Culture and Fine Arts (Kingdom of Cambodia), and the Royal University of Fine Arts (Kingdom of Cambodia). A total of 111 individuals were sorted and analyzed from 57 burial features excavated from excavation unit AB7, a 5 m by 2 m unit near the edge of the cemetery mound. Many of the burials were commingled (in one case the remains of at least five individuals are represented) and lack clear grave cuts. The extensive commingling of the burials is due, in part, to past disturbances of the stratified nature of this densely packed cemetery, often resulting in subsequent interments in the same location^14^. Despite site disturbance, the presence of 33 primary burials, most interred in the same orientation (with head pointing southwest), suggests the excavated portion (approximately 0.03 percent of the mound) was part of a designated communal burial area.

AB M-40/I1680, a Protohistoric Period male individual (dated to 78 − 234 calCE^4^) from the Vat Komnou cemetery, is the same individual as AB40 in Lipson *et al*. (2018)^4^. The individual was sorted from commingled remains of at least five different individuals. The skeletal remains for AB M-40 included cranial and postcranial elements of a male child whose age was estimated to be 5-7 years at the time of death based on clavicular diaphyseal length; the degree of epiphyseal fusion; and dental calcification and eruption. The cranial elements include the right and left parietals, occipital, frontal, right zygomatic, inferior maxillary bones, left temporal, and the petrous portion of the right temporal. The mandible was missing the right ramus. A left clavicular shaft and a thoracic vertebra represent the postcranial remains for this individual. DNA was obtained from the petrous portion of the right temporal bone (for details on DNA extraction and library preparation see Lipson *et al*. 2018^4^).

In the present study, we generated two single-stranded sequencing libraries for this individual (according to a protocol by Gansauge et al. 2017^36^) in addition to 7 analyzed by Lipson et al. (2018)^4^ and changed the strategy for processing BAM files, which increased the coverage for autosomal targets on the 1240K SNP panel^15,16^ from 0.047x to 0.061x and increased the number of usable SNPs from 54,221 to 64,103 1240K sites (see details on all the sequencing libraries in Suppl. Table 2). We merged the new data with published genome-wide data for ancient individuals from Southeast Asia^3,4^ and worldwide present-day populations^24,37-40^ using *PLINK v*.*1*.*90b6*.*10* (https://www.cog-genomics.org/plink/). We kept only autosomal SNPs from the 1240K panel^15,16^. We filtered out ancient individuals who have data for fewer than 50,000 autosomal SNPs. Details of the dataset composition are available in Suppl. Table 3.

### Principal component analysis (PCA)

We performed PCA using *smartpca*^41^ version 16000 from the *EIGENSOFT* package (https://github.com/DReichLab/EIG). We computed principal components for present-day populations from Central, East, Southeast, and South Asia, Andamanese Islands, Siberia, and Europe. We then projected the ancient Southeast Asian individuals onto the principal components using options “lsqproject: YES” and “shrinkmode: YES”.

#### ADMIXTURE

We performed LD-pruning using *PLINK v*.*1*.*90b6*.*10* with options “--indep-pairwise 50 5 0.5” (window size = 50 SNPs, window step = 5 SNPs, r2 threshold = 0.5). The LD-pruned dataset included 523,650 SNPs. We performed *ADMIXTURE* analysis^22^ using *ADMIXTURE v*.*1*.*3* (https://dalexander.github.io/admixture/download.html). We first tested 4 to 12 hypothetical ancestral populations (*K* from 4 to 12) with tenfold cross-validation and five independent algorithm iterations for each *K* value. The CV errors are almost indistinguishable for *K* from 6 to 10 and grow for *K* values above 10 (Suppl. Fig. 2). We subsequently ran up to 30 algorithm iterations for *K* = 6 We chose an iteration with the highest value of the model likelihood as the final result shown in Fig. 3. We visualized the *ADMIXTURE* results using *AncestryPainter* (https://www.picb.ac.cn/PGG/resource.php).

### Outgroup *f*_*3*_-statistics

We used the function “*qp3pop*” from the R package “*ADMIXTOOLS2*”^42^with default settings (https://uqrmaie1.github.io/admixtools/index.html). For outgroup *f*_*3*_-statistics in Fig. 4, we calculated statistics of the form *f*_*3*_(Mbuti; the ancient individual AB M-40/I1680, X), where X are present-day populations from East and Southeast Asia

For the *f*_*3*_-biplot in Fig. 5, we computed statistics *f*_*3*_ (Mbuti; Brahmin Uttar Pradesh, X) and *f*_*3*_ (Mbuti; Dai, X), where X are ancient individuals from Southeast Asia, present-day populations from East, Southeast, and South Asia.

### *qpWave* and *qpAdm*

We used a set of 13 populations as “outgroup” (“right”) populations: 12 diverse worldwide populations (Mbuti, Ami, Japanese, Yi, Onge, Papuan, Pima, Karitiana, Druze, Adygei, Sardinian, and Orcadian) + one of the following South Asian populations (Balochi, Bengali, Brahmin Uttar Pradesh, Brahui, Burusho, Irula, Kalash, Makrani, Mala, Pathan, Punjabi, Rajput, Sindhi, and Vellalar). All the South Asian surrogates were composed of at least three individuals. In total, we had 14 alternative sets of reference populations.

We first performed pairwise *qpWave* modeling using the *qpwave* function from the R package “*ADMIXTOOLS2*”^42^. We considered a pair composed of the target individual AB M-40/I1680 from Cambodia and one of the 13 reference populations as “left populations”. We assigned the remaining 12 reference populations as “right populations”. We used a cut-off *p*-value of 0.05 and tested both “*allsnps = TRUE*” (using all the possible SNPs for each *f*_*4*_-statistic) and “*allsnps = FALSE*” settings (using only SNPs with no missing data at the group level across the “left” and “right” populations). Overall, we tested 182 group pairs with *qpWave* per one target group per one “allsnps” setting.

As none of the *qpWave* models for the individual AB M-40/I1680 was fitting the data, we further tested 2-way admixture models using *qpAdm* with a “rotating” strategy^28^. All possible combinations of two reference populations acted as surrogates for the target. The remaining 11 reference populations were assigned as “right populations”. We defined a model that meets two following criteria as a “plausible model”: 1) *p*-value is higher than 0.05; 2) inferred admixture proportions ± 2 standard errors lie between 0 and 1 for all ancestry components. We tested *qpAdm* models using the *qpadm* function from the R package “*ADMIXTOOLS2*”. We tested both “*allsnps = TRUE*” and “*allsnps = FALSE*” settings. In total, 1,092 models per target per one “allsnps” setting were tested.

## Data Availability

The new genome-wide genotyping data generated in this study will be publicly available when the manuscript is published.

## Acknowledgments

Thanks go to Cambodia’s Ministry of Culture and Fine Arts for permission to collaborate on the Lower Mekong Archaeological Project’s excavations at Vat Komou in the Angkor Borei District. We are grateful to Nadin Rohland and Iñigo Olalde (Harvard University) for generating sequencing libraries and sequencing data for the individual AB M-40/I1680. This work was supported by the Czech Ministry of Education, Youth and Sports: 1) Inter-Excellence program, project #LTAUSA18153; 2) Large Infrastructures for Research, Experimental Development and Innovations project “IT4Innovations National Supercomputing Center – LM2015070”. P.F. was also supported by a subsidy from the Russian federal budget (project No. 075-15-2019-1879 “From paleogenetics to cultural anthropology: a comprehensive interdisciplinary study of the traditions of the peoples of transboundary regions: migration, intercultural interaction and worldview”). The ancient DNA laboratory work was supported by a grant by the John Templeton Foundation (grant 61220) and by the Howard Hughes Medical Institute.

## Author contributions

P.F. and D.R. supervised the study. P.C. designed the study and performed the data analysis. M.P., M.T.S., R.M.I-Q, and R.P. performed archaeological excavations, anthropological analyses of the skeletal material, and sampling for ancient DNA extraction. P.C. and P.F. drafted the manuscript with additional input from all other co-authors.

## Supplementary Information

**Suppl. Fig. 1**. A zoomed-in version of the PCA plot in Fig. 2

**Suppl. Fig. 2**. Cross-validation (CV) errors of the *ADMIXTURE* analysis for different *K* values.

**Suppl. Table 1** Full qpWave and qpAdm results

**Suppl. Table 2** Information about sequencing libraries for the protohistoric period individual AB M-40/I1680

**Suppl. Table 3** Dataset composition

## References

1. Eberhard, D., Simons, G. F. & Fennig, C. D. Ethnologue. Languages of Asia, Twenty-third edition. (SIL International, Global Publishing, 2020).

2. O’Connor, S. & Bulbeck, D. Homo Sapiens Societies in Indonesia and South-Eastern Asia. in The Oxford Handbook of the Archaeology and Anthropology of Hunter-Gatherers (eds. Cummings, V., Jordan, P. & Zvelebil, M.) 346–367 (Oxford University Press, 2014).

3. McColl, H. et al. The prehistoric peopling of Southeast Asia. Science 361, 88–92 (2018).

4. Lipson, M. et al. Ancient genomes document multiple waves of migration in Southeast Asian prehistory. Science 361, 92–95 (2018).

5. Cœdès, G. The Indianized states of Southeast Asia. (University of Hawaii Press, 1968).

6. Bellina, B. Southeast Asian Evidence for Early Maritime Silk Road Exchange and Trade-Related Polities. in The Oxford Handbook of Early Southeast Asia (eds. Higham, C. F. W. & Kim, N. C.) 457–500 (Oxford University Press, 2022).

7. Stark, M. T. Landscapes, Linkages, and Luminescence: First-Millennium CE Environmental and Social Change in Mainland Southeast Asia. in Primary Sources and Asian Pasts (eds. Bisschop, P. C. & Cecil, E. A.) 184–219 (De Gruyter, 2020).

8. Mörseburg, A. et al. Multi-layered population structure in Island Southeast Asians. Eur. J. Hum. Genet. 24, 1605–1611 (2016).

9. Kutanan, W. et al. Reconstructing the Human Genetic History of Mainland Southeast Asia: Insights from Genome-Wide Data from Thailand and Laos. Mol. Biol. Evol. 38, 3459–3477 (2021).

10. Changmai, P. et al. Indian genetic heritage in Southeast Asian populations. PLOS Genet. 18, e1010036 (2022).

11. Hellenthal, G. et al. A genetic atlas of human admixture history. Science 343, 747–751 (2014).

12. Stark, M. T. & Sovath, B. Recent research on emergent complexity in Cambodia’s Mekong. Bull. Indo-Pacific Prehistory Assoc. 21, 85–98 (2001)

13. Stark, M. T. Some Preliminary Results of the 1999-2000 Archaeological Field Investigations at Angkor Borei, Takeo Province. Udaya J. Khmer Stud. 2, 19–35 (2001).

14. Ikehara-Quebral, R. M. An assessment of health in Early Historic (200 BC to AD 200) inhabitants of Vat Komnou, Angkor Borei, southern Cambodia: A bioarchaeological perspective, Ph.D. dissertation (University of Hawai‘i at Mānoa, 2010).

15. Fu, Q. et al. An early modern human from Romania with a recent Neanderthal ancestor. Nature 524, 216–219 (2015).

16. Rohland, N. et al. Three Reagents for in-Solution Enrichment of Ancient Human DNA at More than a Million SNPs. bioRxiv 2022.01.13.476259 Preprint at https://www.biorxiv.org/content/10.1101/2022.01.13.476259v1 (2022).

17. Reich, D., Thangaraj, K., Patterson, N., Price, A. L. & Singh, L. Reconstructing Indian population history. Nature 461, 489–494 (2009).

18. Chaubey, G. et al. Population Genetic Structure in Indian Austroasiatic Speakers: The Role of Landscape Barriers and Sex-Specific Admixture. Mol. Biol. Evol. 28, 1013–1024 (2011).

19. Nakatsuka, N. et al. The promise of discovering population-specific disease-associated genes in South Asia. Nat. Genet. 49, 1403–1407 (2017).

20. Tätte, K. et al. The genetic legacy of continental scale admixture in Indian Austroasiatic speakers. Sci. Rep. 9, 3818 (2019).

21. Jeong, C. et al. The genetic history of admixture across inner Eurasia. Nat. Ecol. Evol. 3, 966–976 (2019).

22. Alexander, D. H., Novembre, J. & Lange, K. Fast model-based estimation of ancestry in unrelated individuals. Genome Res. 19, 1655–1664 (2009).

23. Mondal, M. et al. Genomic analysis of Andamanese provides insights into ancient human migration into Asia and adaptation. Nat. Genet. 48, 1066–1070 (2016).

24. Bergström, A. et al. Insights into human genetic variation and population history from 929 diverse genomes. Science 367, eaay5012 (2020).

25. Jeong, C. et al. Bronze Age population dynamics and the rise of dairy pastoralism on the eastern Eurasian steppe. Proc. Natl. Acad. Sci. U. S. A. 115, E11248–E11255 (2018).

26. Zhang, C. et al. Differentiated demographic histories and local adaptations between Sherpas and Tibetans. Genome Biol. 18, 115 (2017).

27. Haak, W. et al. Massive migration from the steppe was a source for Indo-European languages in Europe. Nature 522, 207–211 (2015).

28. Harney, É., Patterson, N., Reich, D. & Wakeley, J. Assessing the performance of qpAdm: a statistical tool for studying population admixture. Genetics 217, iyaa045 (2021).

29. Richard Scott, G. & Turner, C. G. The Anthropology of Modern Human Teeth Dental Morphology and its Variation in Recent Human Populations. (Cambridge University Press, 1997)

30. Shewan, L. et al. Resource utilisation and regional interaction in protohistoric Cambodia – The evidence from Angkor Borei. J. Archaeol. Sci. Reports 31, 102289 (2020).

31. Manguin, P.-Y. & Stark, M. T. Mainland Southeast Asia’s Earliest Kingdoms and the Case of “Funan”. in The Oxford Handbook of Early Southeast Asia (eds. Higham, C. F. W. & Kim, N. C.) 637–659 (Oxford University Press, 2022).

32. Carter, A. K., Dussubieux, L., Stark, M. T. & Gilg, H. A. Angkor Borei and Protohistoric Trade Networks: A View from the Glass and Stone Bead Assemblage. Asian Perspect. 60, 32–70 (2020).

33. Castillo, C. C. et al. Rice, beans and trade crops on the early maritime Silk Route in Southeast Asia. Antiquity 90, 1255–1269 (2016).

34. Bellina, B. Maritime Silk Roads’ Ornament Industries: Socio-political Practices and Cultural Transfers in the South China Sea. Cambridge Archaeol. J. 24, 345–377 (2014).

35. Zakharov, A. The Angkor Borei Inscription K. 557/600 from Cambodia: An English translation and commentary. Vostok. Afro-aziatskie Obs. Istor. i Sovrem. 66–80 (2019).

36. Gansauge, M. T. et al. Single-stranded DNA library preparation from highly degraded DNA using T4 DNA ligase. Nucleic Acids Res. 45, e79–e79 (2017).

37. Raghavan, M. et al. Upper Palaeolithic Siberian genome reveals dual ancestry of Native Americans. Nature 505, 87–91 (2014).

38. Raghavan, M. et al. Genomic evidence for the Pleistocene and recent population history of Native Americans. Science 349. aab3884 (2015).

39. Mallick, S. et al. The Simons Genome Diversity Project: 300 genomes from 142 diverse populations. Nature 538, 201–206 (2016).

40. de Barros Damgaard, P. et al. The first horse herders and the impact of early Bronze Age steppe expansions into Asia. Science 360, eaar7711 (2018).

41. Patterson, N., Price, A. L. & Reich, D. Population structure and eigenanalysis. PLoS Genet. 2, e190 (2006).

42. Maier, R., Flegontov, P., Flegontova, O., Changmai, P. & Reich, D. On the limits of fitting complex models of population history to genetic data. bioRxiv 2022.05.08.491072 Preprint at https://www.biorxiv.org/content/10.1101/2022.05.08.491072v2 (2022).

