## Supplementary figures and images for "Ancient DNA from Protohistoric Period Cambodia indicates that South Asians admixed with local populations as early as 1^st^-3^rd^ centuries CE"

### Supplemental Figure 1

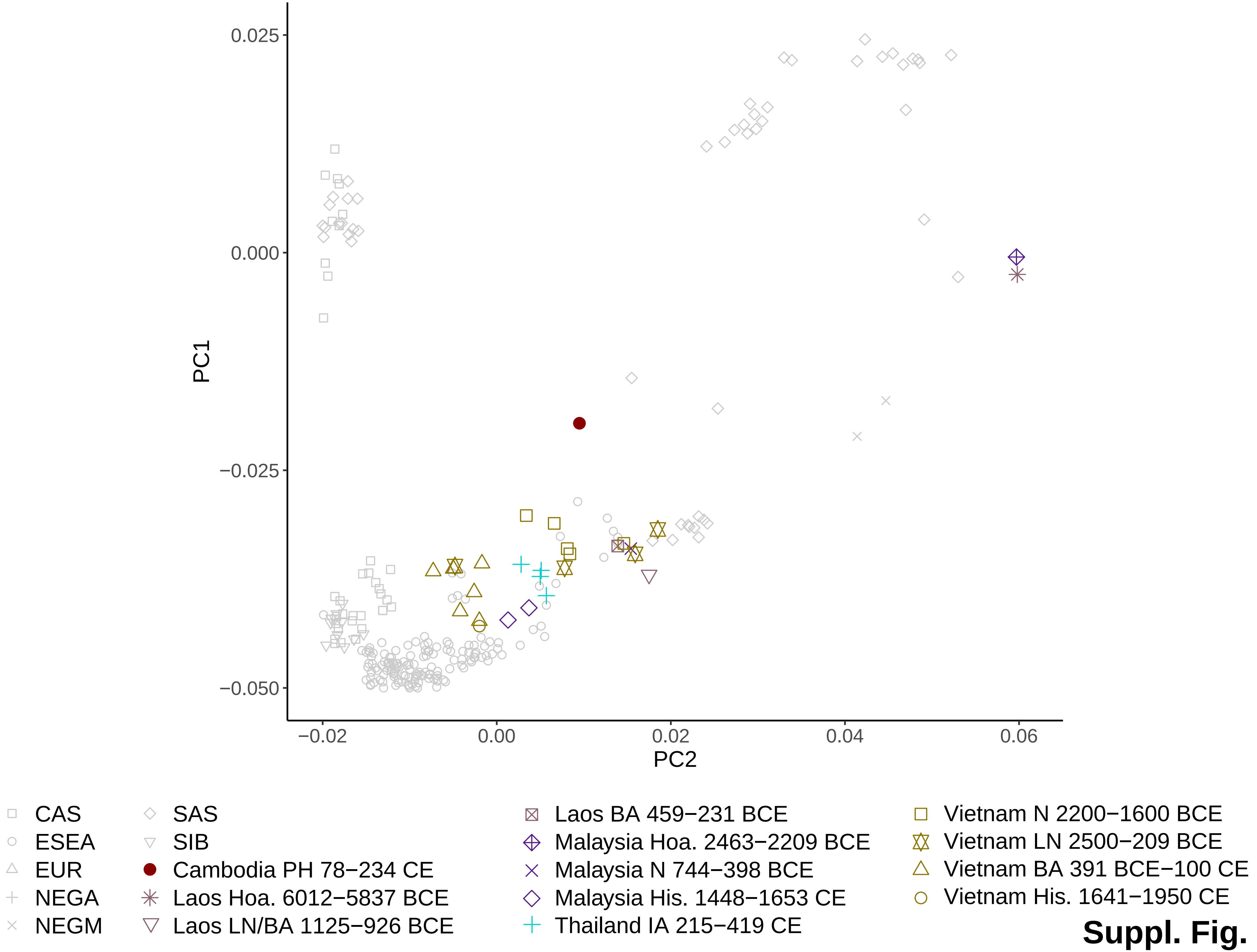

### Supplemental Figure 2

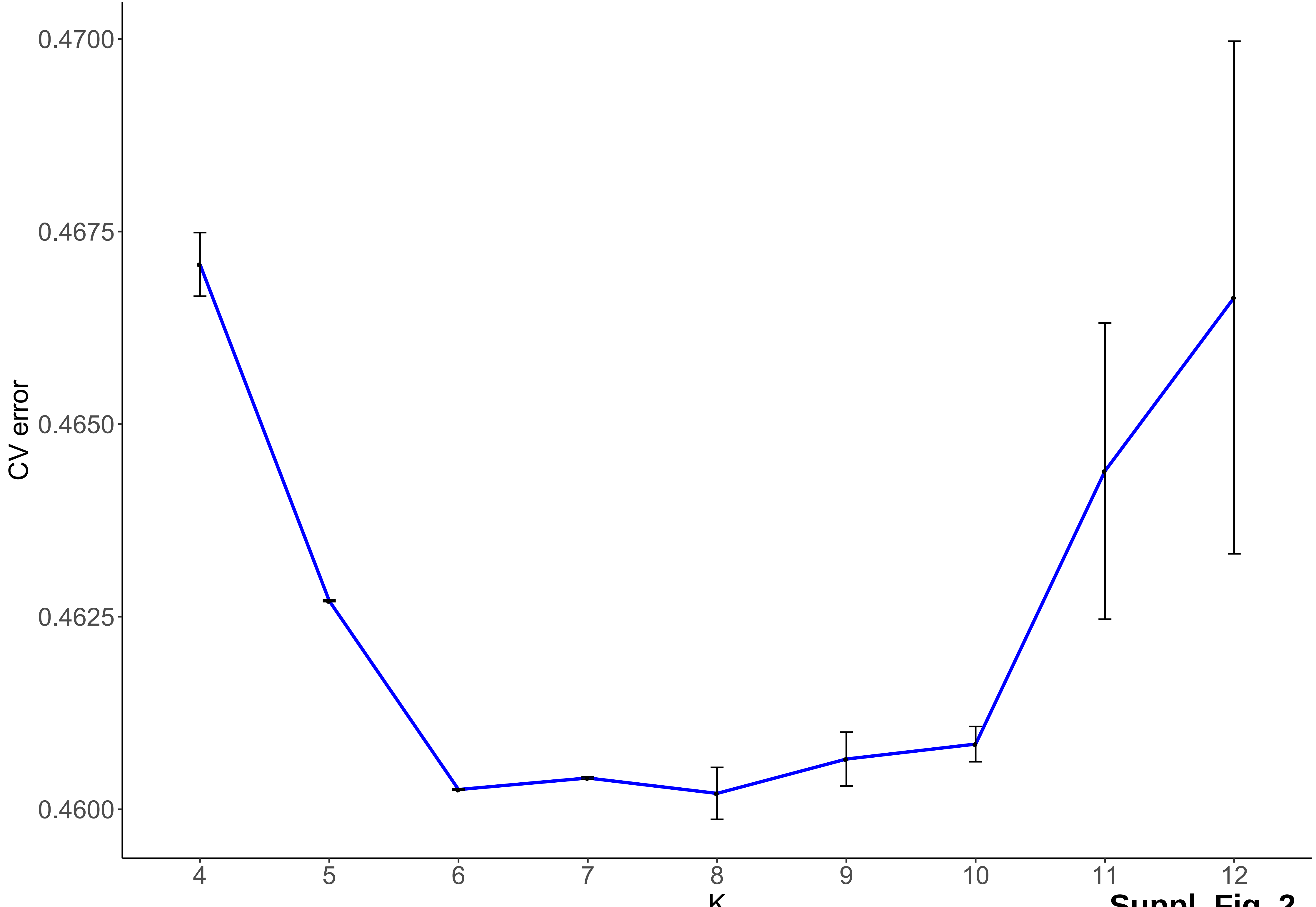

Suppl. Fig. 2
